# Spatial control of Keratin 8 phosphorylation by Aurora B facilitates cytokinesis in cancer cells of epithelial origin

**DOI:** 10.1101/2023.01.22.525045

**Authors:** Busra Harmanda, Halenur Ayaydin, Xenia Waide, Mohammad H Qureshi, Venkatesha Basrur, Alexey I Nesvizhskii, Timothy J Mitchison, Nurhan Ozlu

## Abstract

Keratins assemble into mechanically resilient polymers that physically stabilize epithelial cells. When epithelial cells divide, keratin polymers must be severed to allow cell separation during cytokinesis. Phosphorylation has been implicated in this process, but how keratins are regulated during cell division is not understood. Aurora-B kinase, which is part of the chromosome passenger complex (CPC) accumulates at the cell center during cytokinesis and has been implicated in regulating intermediate filaments. We mapped six Aurora B kinase sites in Keratin 8. Phosphorylation of Keratin 8 at S34 occurred specifically at the cleavage furrow and persisted at the midzone until the completion of cytokinesis. Inhibition of Aurora B or expression of a non-phosphorylatable Keratin 8 mutant impaired keratin disassembly at the cleavage furrow. We propose that Aurora B–mediated phosphorylation promotes localized keratin filament disassembly at the cleavage furrow, allowing spatially regulated disassembly during cytokinesis. Aurora B binds to keratin filaments, and its localization to midzones was reduced in Keratin 8 knockout cells, showing that Keratin 8 facilitates Aurora B targeting during cytokinesis. This suggests a positive feedback cycle whereby Keratin 8 promotes midzone-localization of Aurora B and in turn, is locally disassembled by its kinase activity. This cycle is required for successful furrow ingression and completion of cell division in cancer cells of epithelial origin and might provide a target for solid tumor treatment.

## Introduction

Keratin filaments are a class of intermediate filaments that are abundant in epithelial cells, where they provide mechanical integrity [1, 2]. Keratin proteins are expressed in pairs that heterodimerize to generate polymerizing subunits, with different cell types expressing characteristic pairs [3]. Keratin 8/18 subunits are simple epithelial keratins typically expressed in proliferating epithelial progenitor cells as well as many epithelial cancers and cancer-derived cell lines [4] and are thus of particular relevance when considering the roles of keratins in cell division in epithelial tissues and cancers of epithelial origin (carcinomas).

The primary functional concern in the biology of keratins during cell division is the need to remove them, in whole or part, to allow the successful execution of mitosis and cytokinesis. Keratin filament networks, which span the whole cytoplasm and connect between cells at desmosomes [5], have the potential to physically impede mitotic spindle assembly and cleavage furrow ingression. Early studies suggested the Keratin 8/18 network globally disassembles into non-filamentous aggregates at the onset of mitosis [6]. Later work showed that in some epithelial cell types, keratin disassembly is spatially organized to generate a filament-free cage in which the mitotic spindle assembles [7]. Temporal and spatial control of intermediate filament organization during cell division is exerted by kinases that phosphorylate subunits and typically drive the monomer-filament system towards disassembly [8]. Control of vimentin polymerization by phosphorylation during cell division has been studied in depth, and it is known that Aurora-B kinase has an important regulatory role [9, 10]. Control of keratin organization during cell division has been much less studied, despite the importance of keratins in tissue biology and epithelial cancers. Keratins are heavily phosphorylated and their phosphorylation levels are enhanced during mitosis [11]. Keratins are probably phosphorylated by multiple kinases during mitosis. Keratin 5/14 were reported to be phosphorylated by CDK1, Rho-kinase and Aurora B kinases during cell division [8].

Most studies portray intermediate filaments as potential blocks to cell division that must be removed. However, emerging studies showed that beyond their classical role in providing mechanical support, keratins have been implicated in various regulatory processes including cell signaling, cell growth, cell differentiation, apoptosis and stress response [12–14]. Similarly, vimentin filaments have been shown to promote spatially organized signaling during cell migration [15, 16]. Vimentin is thought to serve as a scaffold that promotes the local activity of ERK kinase [17]. It is possible that K8/18 filament contribute to the spatial organization of cell division in epithelial progenitors and cancers, for example by helping scaffold mitotic kinases, but this hypothesis has to be systematically addressed.

Aurora B kinase acts in cell division as part of the CPC. During metaphase and anaphase CPC localizes to centromeres and the surface of chromosomes where it regulates chromosome condensation and helps correct errors in kinetochore-microtubule attachments [18]. At the onset of cytokinesis, it translocates to the spindle midzone where it promotes assembly and ingression of the cleavage furrow. Midzone-localized Aurora B also generates a phosphorylation gradient that helps to resolve segregation errors in between segregating chromosomes [19, 20] and delays abscission when unsegregated chromosomes are trapped at the cleavage site [21, 22].

In our previous phospho-proteomic studies, we found that phosphorylations on Keratin 8 and 18 were diminished after Aurora B kinase inhibitor treatment [23–25]. In Xenopus laevis eggs, a model system for epithelial cell division, keratins are globally disassembled during mitosis, presumably by CDK1 activity [26, 27], and locally disassembled between microtubule asters during interphase by Aurora B kinase activity [28].

In this study, by taking mass spectrometry-based *in vitro* kinase assay, we systematically mapped Aurora B-dependent phosphorylations of Keratin 8. Aurora B kinase interacts with Keratin 8 in a cell-cycle dependent manner. Aurora B-dependent Keratin 8 phosphorylation (K8 phosphoS34) decorates the cleavage furrow and facilitates Keratin 8 disintegration during furrow ingression. Ectopic expression of non-phosphorylatable Keratin 8 dramatically increases cleavage furrow regression. We propose that the interaction between Aurora B kinase and Keratin 8 has mutual effects: it helps translocation of Aurora B to the midzone and promotes Keratin disintegration during cleavage furrow ingression in epithelial cells and epithelial malignancies.

## Results

### Aurora-B phosphorylates Keratin 8 during cell division

Our previous proteomic analysis suggested that Keratin 8 is phosphorylated by Aurora B kinase during cell division [25]. To test this, we performed an *in vitro* kinase assay followed by mass spectrometry analysis. We expressed the Keratin 8-GST fusion protein in bacteria and purified it using glutathione beads. Keratin 8-GST and active Aurora B complexed with a fragment of INCENP were incubated in a kinase reaction buffer in the presence of ATP. A parallel reaction that has only Keratin 8-GST but lacks Aurora B complex is performed as a control **(Figure 1A)**. The Aurora B-dependent Keratin phosphorylations are quantified by monitoring Keratin 8 phosphopeptides and their non-phosphorylated counterparts by PRM (Parallel Reaction Monitoring) experiments using Mass Spectrometry. This targeted proteomics approach is well-suited for phosphorylation studies, as it enhances both sensitivity and reproducibility in site-specific analysis. By selectively monitoring predefined precursor–fragment ion pairs of the phosphopeptide of interest, PRM minimizes interference from co-eluting peptides and enables precise, reproducible quantification across experimental conditions [29]. Keratin 8 S34 phospho and nonphospho-forms are quantified in both − /+ Aurora B kinase set-up **(Figure 1B, Supplementary S1A-F)**. By this approach, we map the Aurora B kinase-dependent Keratin 8 phosphosites *in vitro*. As a result, S34, S37, S124, S330, S404 and S475 Keratin 8 residues were phosphorylated in the presence of Aurora B **(Figure 1C, Supplementary S1A-F)**. In a previous global phosphoproteome study, we analyzed cell cycle-dependent phosphorylations and found that the phosphosites S13, S34, S35, S36, S37, S39, S43, S44, S104, and S258 of Keratin 8 are up-regulated during mitosis or cytokinesis [24].

**Figure 1.**
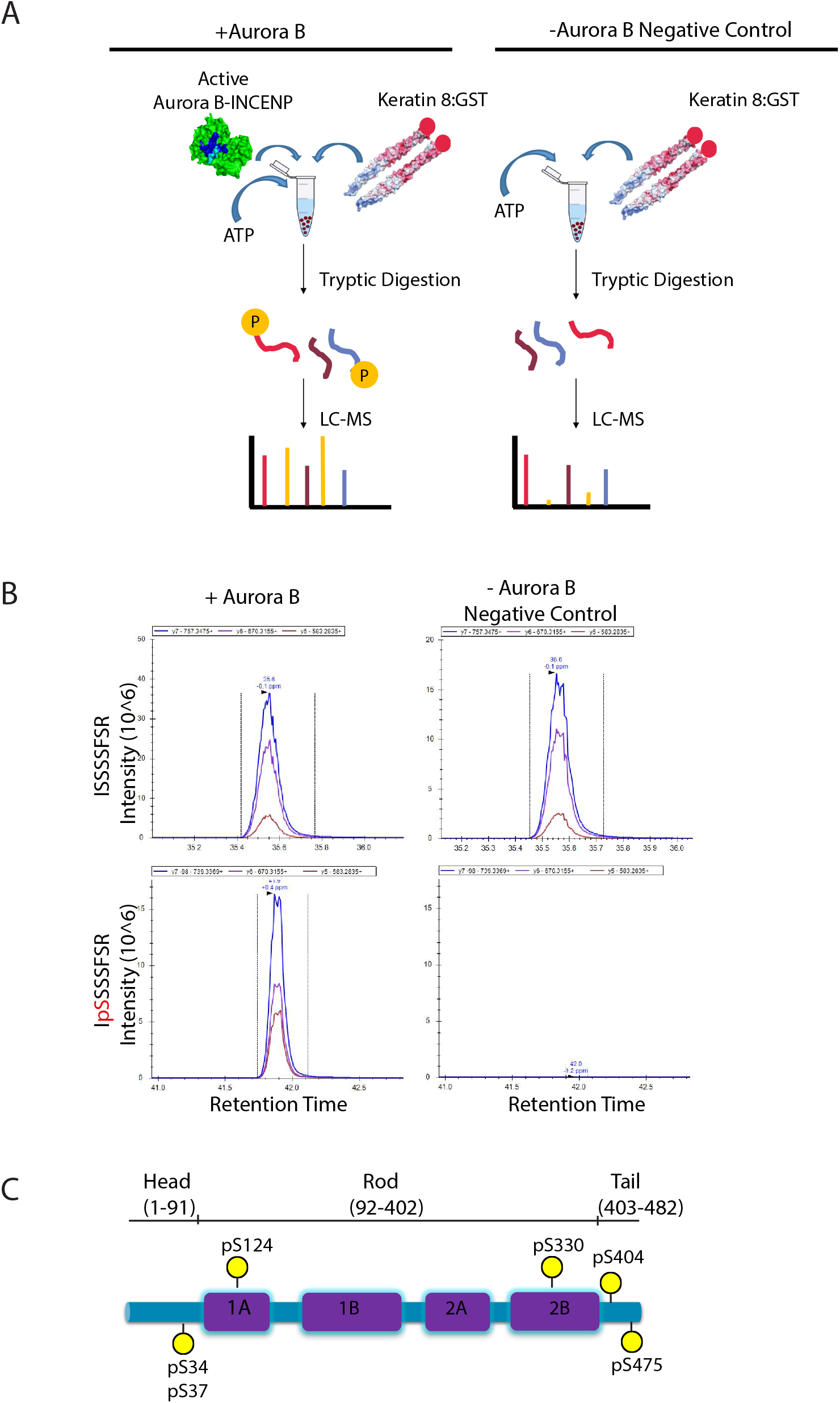
Mapping Aurora B kinase-dependent phosphorylation sites in Keratin 8 by *in vitro* kinase assay. **A**. Cartoon illustration of the *in vitro* kinase assay workflow for identifying Aurora B phosphorylation sites on Keratin 8. **B**. Parallel Reaction Monitoring-Mass Spectrometry (PRM-MS) analysis of the *in vitro* phosphorylation of S34 Keratin 8 by Aurora B (n=1). The peak area quantification of the nonphospho (ISSSSFSR) and phosphoS34 (IpSSSSFSR) of Keratin 8 with Aurora B (+Aurora B) and without Aurora B (-Aurora B) conditions. Each graph shows one peptide and the colored peaks show the transitions of the non-phosphopeptide (top) and phosphopeptide (bottom). **C**. The map of Aurora B-dependent K8 phosphorylation sites detected by the *in vitro* kinase assay (n=1). Yellow, phosphorylated Serine residues on the head, rod or tail domains of Keratin 8.

### Aurora B-dependent Keratin 8 phosphorylation is required for cytokinesis

To analyze the role of Aurora B-dependent phosphorylation of Keratin 8 (K8), we decided to focus on S34-37 which is a highly phosphorylated region at the head domain **(Figure 1C)**. In agreement with the in vitro kinase assay, our previous study also identified S34 and S37 Keratin as cytokinesis selective phosphorylation sites whose levels were diminished when cells were treated with an Aurora inhibitor, VX680 [24]. To test the role of these phosphorylation sites in cell division, we created K8 S34-35A, S36-37A, S34-37A phosphomutants and S34-35D, S36-37D, S34-37D phosphomimetic mutants fused to GFP and expressed them in Keratin 8 Knockout (K8 KO) cells. K8 KO HeLa cells were generated using the CRISPR/Cas9 system (**Supplementary Figure S2**). The knockout of Keratin 8 in these K8 KO cells was confirmed through Western blotting analyses **(Supplementary Figure S2A)**. Non-targeting guide RNA-expressing cells served as controls in these assays. We then quantified multinucleation, which provides a simple readout of cytokinesis failure (**Supplementary Figure S2B)**. Western blotting analysis showed that total protein levels of all mutated K8 are comparable **(Supplementary Figure S2C)**. We observed a significant increase in multinucleation of 4xA (S34-37A) K8-GFP cells, but not in 2xA mutants (S34-S35, S36-S37). In contrast to the 4xA mutation, 4xD (S34-37D) did not cause a significant increase in multinucleation **(Figure 2A, Supplementary Figure S2B)**. This result suggests that non-phosphorylatable Keratin 8 perturbs cytokinesis, and phosphorylation of S34-37 has an impact on cytokinesis.

**Figure 2.**
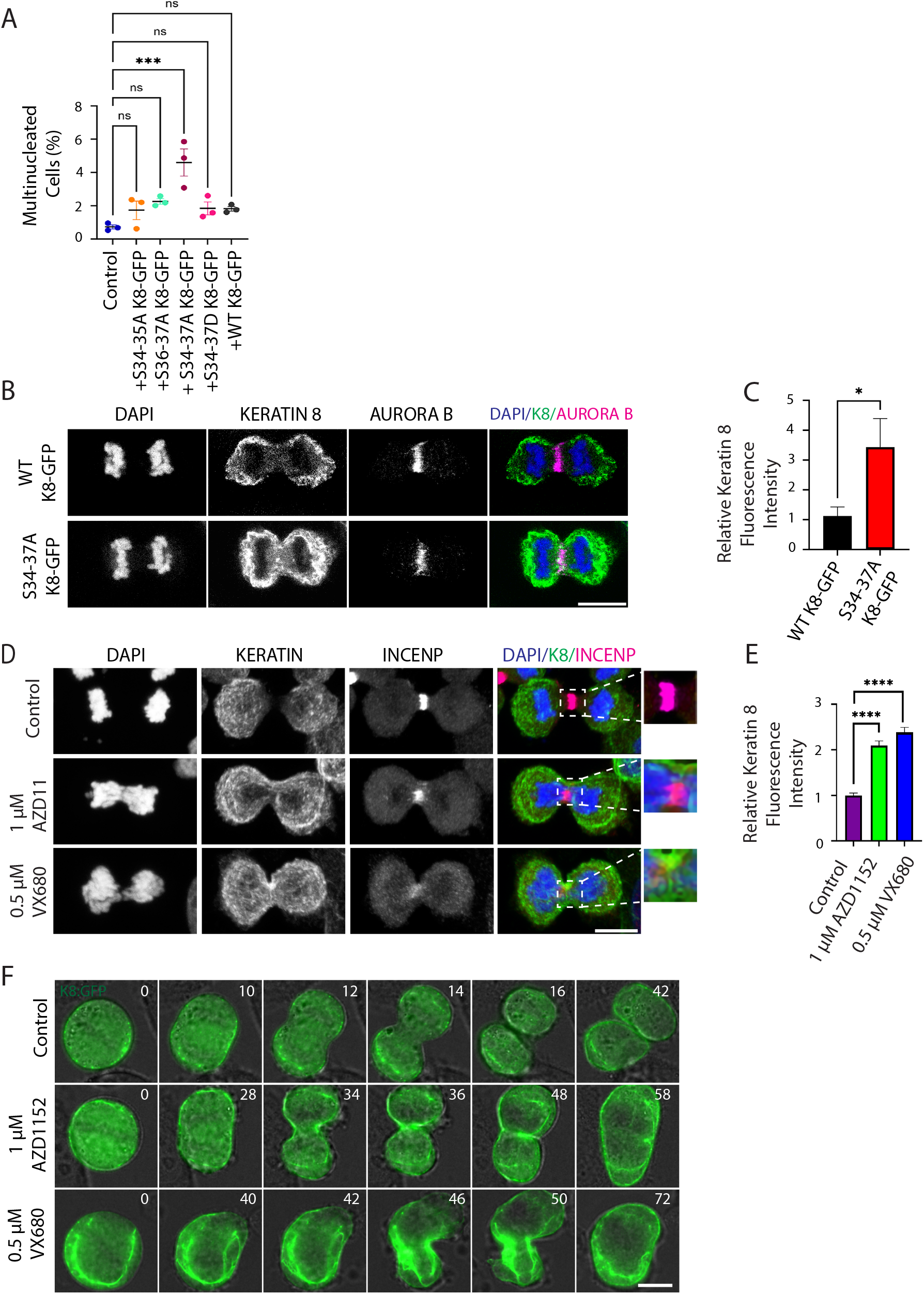
Non-phosphorylatable Keratin 8 mutation causes persistent Keratin 8 bundles during cleavage furrow ingress. **A**. Quantification of the percentage of multinucleated cells in control and K8 Knockout HeLa cells expressing WT K8-GFP (n=1477), S34-35A K8-GFP (n=1568), S36-37A K8-GFP (n=796), S34-37A K8-GFP (n=1450), and S34-37D K8-GFP (n=1371) from three independent experiments. Statistical analysis was performed using one-way ANOVA with the Brown-Forsythe test. Error bars represent the standard error of the mean (SEM). *** p < 0.001; ns, not significant. **B**. Representative images of K8 Knockout HeLa cells expressing WT K8-GFP (n=20) and S34-37A K8-GFP (n=20) during cytokinesis from three independent experiments. Images show GFP-tagged WT and S34-37A mutant K8 proteins (green), Aurora B (magenta), and DNA staining (DAPI, blue). Scale bar, 10 μm. **C**. Quantification of WT K8-GFP and S34-37A K8-GFP localization at the cleavage furrow in HeLa cells. K8-GFP fluorescence intensities at the cleavage furrow were measured and normalized to cytosolic K8-GFP intensities. n = 20 cells per group from three independent experiments. Statistical analysis was performed using an unpaired two-tailed t-test. Data are presented as mean ± SEM. * p < 0.05. **D**. Immunofluorescence staining for Keratin 8 (green), INCENP (magenta) and DAPI (blue) in control (n=65) and HeLa cells treated with 1 µM AZD1152 (n=48) or 0.5 µM VX680 (n=57) after thirty minutes of nocodazole release. Insets show a magnified view of the midzone region. Scale bar, 10 µm. **E**. Quantification of Keratin 8 localization at the cleavage furrow in control (n=65), 1 µM AZD1152 (n=48) or 0.5 µM VX680 (n=57) treated HeLa cells. One-way ANOVA with Dunnett’s post hoc test was performed. **** p<0.001. Data represent mean ± SEM. **F**. Representative images from live imaging of K8 Knockout cells expressing WT K8-GFP that are treated with DMSO (control n=5) or Aurora B inhibitors (1 µM AZD1152 n=5 or 0.5 µM VX680 n=5) (Videos S1-3). Scale bars, 10 µm.

To further investigate the impact of Aurora B-dependent phosphorylation on Keratin 8, we closely examined Wild-Type (WT) and phospho-mutant (4XA) K8-GFP-expressing cells during cytokinesis. In WT K8-GFP-expressing HeLa cells, Keratin 8 disappeared at the cleavage furrow of cytokinesis cells. Interestingly, 4XA K8-GFP-expressing cells exhibited residual Keratin 8 at the cleavage furrow **(Figure 2B)**. WT K8 was cleared from a region a few microns wide around the ingression furrow, while 4XA K8 intensity was detectable in bundles at the center of the cell. Quantification of WT and mutant K8-expressing cells supported significantly more Keratin 8 bundles at the cleavage furrow in 4XA K8-GFP-expressing cells than WT ones (**Figure 2C).**

If Aurora B-dependent phosphorylation of Keratin 8 is required for the dissolving of Keratin 8 filaments at the cleavage furrow, then Aurora B inhibition should affect the Keratin 8 dynamics and distribution during cytokinesis. To test this, we synchronized HeLa cells at mitosis with nocodazole and then released them to cytokinesis. Thirty minutes after the release, we treated the cells with Aurora B inhibitors (AZD1152 and VX680) for a further thirty minutes. We performed immunostaining in control and AZD1152/VX680 treated cells using anti-Keratin 8 and anti-INCENP antibodies **(Figure 2D)**. Consistent with the results above, Keratin 8 persisted and, in some cells, formed large bundles at the cleavage furrow upon Aurora B inhibition, whereas Keratin 8 disintegrated and was not visible at the midzone in control cells (**Figure 2D)**. Quantification of multiple control and AZD1152 or VX680 treated cells revealed more than two-fold increase in the Keratin intensity at the cleavage furrow (**Figure 2E)**. Live imaging of AZD1152 or VX680-treated K8-GFP-expressing cells also confirmed those results (**Figure 2F, Videos S1-3**). Strikingly, we observed large bundles of Keratin filaments in Aurora inhibitor-treated cells. These results support that phosphorylation of Keratin 8 by Aurora B kinase is required for furrow ingression. A limiting factor in the use of small-molecule Aurora inhibitors is off-target or pleiotropic effects. VX680 is a pan-Aurora inhibitor with the highest potency toward Aurora A, while AZD1152 is more selective for Aurora B but can also inhibit Aurora A and C at higher concentrations [25, 30]. However, Aurora A is largely degraded at cytokinesis; the likelihood that VX680 effects reflect Aurora A inhibition at this stage is reduced. Aurora B–dependent phosphorylation of K8 by *in vitro* kinase assay **(Figure 1)** and dual-inhibitor strategy strengthen attribution to Aurora B, but off-target activities and pleiotropic consequences at the signaling network level cannot be fully excluded.

### Phosphomimetic mutant of Keratin 8 leads to speckled Keratins

To investigate how K8 phosphorylation regulates filament organization, we examined the cellular distribution of the phosphomutant (4xA) and phosphomimetic (4xD) mutant K8-GFP in K8 KO cells throughout cell division using confocal microscopy **(Figure 3A)**. Live cell imaging revealed that the phosphomimetic mutant of K8 has rather different morphology **(Figure 3A, Videos S4-6)**. In agreement with our previous observations, 4XA K8-GFP often accumulated at the cleavage furrow during cytokinesis. In contrast, we did not observe keratin bundles in 4xD K8-GFP; they rather formed speckled keratins throughout mitosis and cytokinesis **(Figure 3A)**. Next, we analyzed speckled keratins in WT, S34-35A, S34-35D, S34-37A, and S34-37D K8-GFP-expressing cells fixed in paraformaldehyde **(Figure 3B)**. Quantification of both S34-35D and S34-37D K8-GFP-expressing cells revealed a significantly higher percentage of cells (> 90%) with speckled keratins in cytokinesis, with no statistically significant differences between these two groups **(Figure 3C)**. A previous study reported that phosphomimetic Keratin 5mutants display similar speckle-like structures and accumulate as soluble heterotetramers [31]. We suggest that the phosphorylation of S34-37 enhances keratin solubility and promotes the disruption of keratin bundles

**Figure 3.**
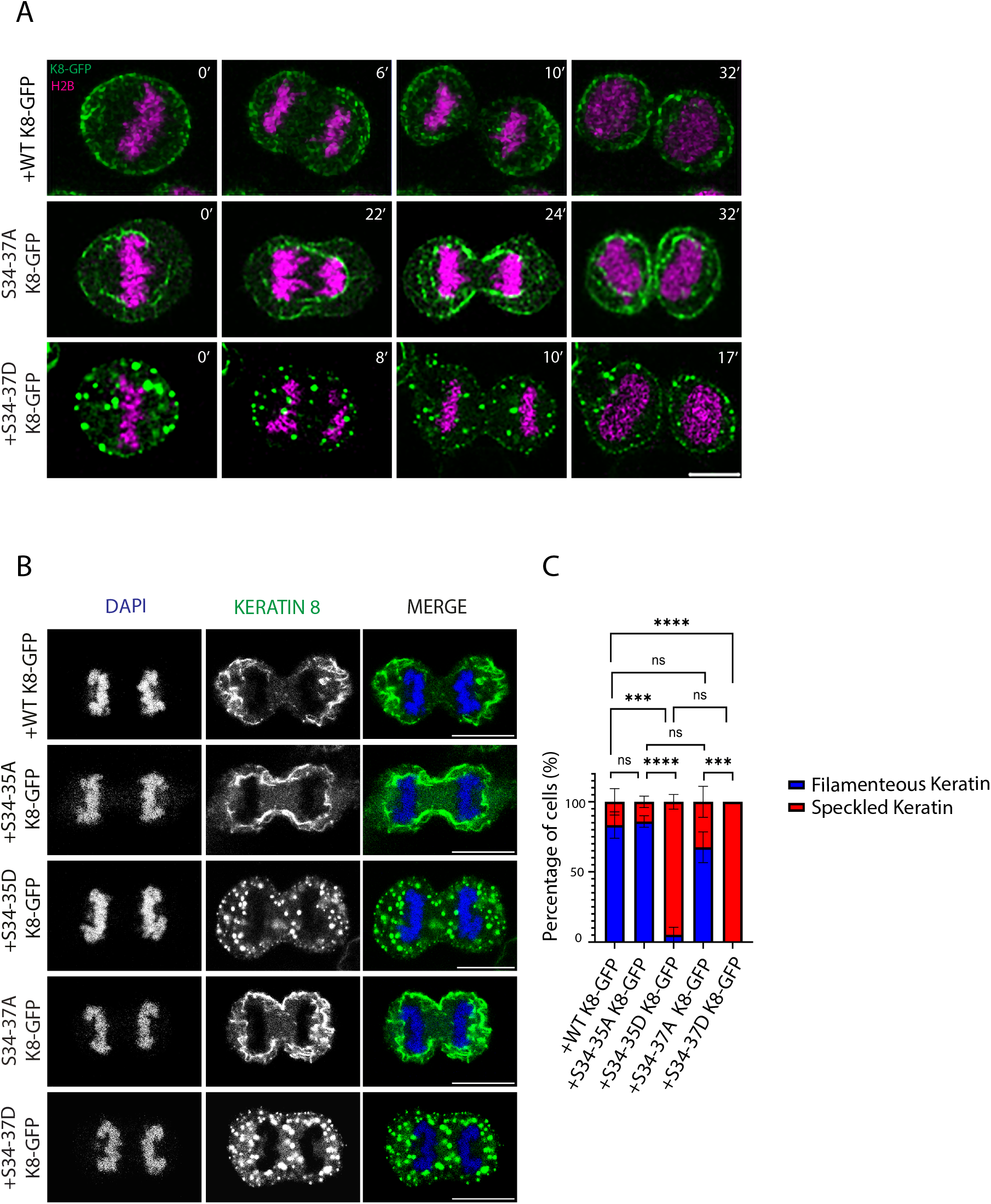
Phosphomimetic mutant of Keratin 8 leads to speckled keratin distribution. **A**. Representative images showing the localization of Keratin 8 phosphomutants during cell division. Still images from time-lapse movies of K8 Knockout HeLa cells expressing mCherry-H2B and either WT K8-GFP, S34-37A K8-GFP, or S34-37D K8-GFP are shown (Videos S4-6) (n=3). The nucleus is shown in magenta and K8-GFP in green. Scale bar, 10 µm. **B**. Representative images showing the localization of Keratin 8 in K8 Knockout HeLa cells expressing WT K8-GFP (n=153), S34-35A K8-GFP (n=24), S34-35D K8-GFP (n=39), S34-37A K8-GFP (n=127), and S34-37D K8-GFP (n=131). HeLa cells were immunostained against Keratin 8 (green), and DNA (DAPI, blue). Scale bars, 10 µm. **C**. Quantification of speckled Keratin structures in dividing K8 Knockout HeLa cells expressing WT K8-GFP (n=153), S34-35A K8-GFP (n=24), S34-35D K8-GFP (n=39), S34-37A K8-GFP (n=127), or S34-37D K8-GFP (n=131) were analyzed per group from two independent experiments. Statistical analysis was performed using two-way ANOVA. ***p=0.0004 ****p < 0.0001; ns, not significant. Data represent mean ± SEM.

### Aurora B-dependent Phosphorylation of Keratin 8 decorates the cleavage furrow

Within the S34–S37 region, S34 aligns with the Aurora B consensus motif, and our *in vitro* kinase assay confirmed that its phosphorylation is dependent on Aurora B activity. Therefore, to examine the Aurora B-dependent spatial and temporal regulation of K8 phosphorylation during division, we raised custom phospho-specific antibodies against phospho S34 K8. We initially confirmed the specificity of the antibody by immunostaining K8 KO cells (**Supplementary Figure S3**). To image the subcellular localization of phosphorylated Keratin 8, unsynchronized HeLa cells were fixed with paraformaldehyde and immunostained using anti-K8 and anti-Phospho S34 K8 antibodies. Although Keratin 8 decorated the inside of cells throughout the cell cycle, phospho S34 K8 was only detectable at the onset of anaphase. As chromosomes segregate, it is strongly localized at the contractile ring while the cleavage furrow ingresses and persists at the neighboring cortical part of newly divided daughter cells during the process of abscission (**Figure 4A**). To confirm the phosphorylation of S34 K8 by Aurora B kinase in vivo, we treated cells with Aurora B kinase inhibitors AZD1152 or VX680 and examined Phospho S34 K8 levels. Notably, Aurora B kinase activity inhibition abolished the contractile ring localization of S34 Keratin 8 phosphorylation (**Figure 4B**). Phosphorylated S34 Keratin 8 absolutely disappeared in both anaphase and telophase cells upon Aurora B inhibition (**Figure 4C**). To test whether this phosphorylation pattern is conserved in Keratin-expressing cells, in addition to HeLa cells, we also tested the localization of Phospho S34 K8 in an epithelial cell line, MCF7. Similar to the expression pattern in HeLa cells, Phospho S34 K8 was only detectable at the onset of anaphase and it was confined to the cleavage furrow until the end of cytokinesis (**Supplementary Figure S4A**). When MCF7 cells were treated with Aurora B inhibitors, phospho S34 staining was completely abolished (**Supplementary Figure S4B**). These results support the conservation of cytokinesis-specific and Aurora B-dependent phosphorylation of Keratin 8 in all Keratin-expressing cells.

**Figure 4.**
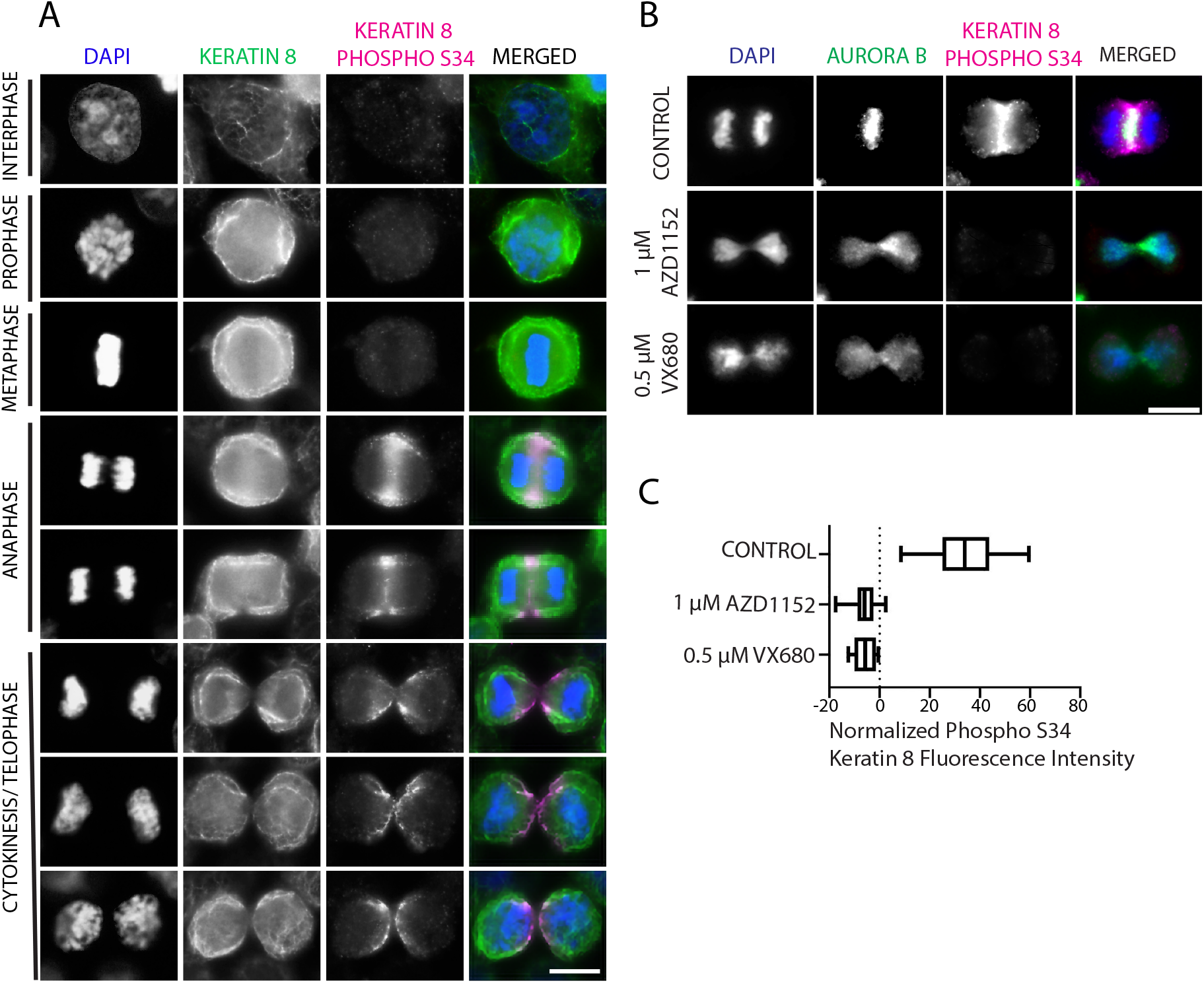
Aurora B-dependent Keratin 8 S34 phosphorylation decorates the cleavage furrow. **A**. Representative images of Keratin 8 S34 phosphorylation at the cleavage furrow. Interphase (n=20), prophase (n=10), metaphase (n=10), anaphase (n=10) and telophase/cytokinesis (n=15) HeLa cells were immunostained against Keratin 8 (green), Keratin 8 phospho S34 (magenta) and DNA (DAPI, blue). Scale bar, 10 µm. **B**. Representative images of HeLa cells after Aurora B inhibition. Control (n=11), 1 µM AZD1152 (n=9) or 0.5 µM VX680 (n=8) treated cells were immunostained against Aurora B (green), Keratin 8 phospho S34 (magenta) and DNA (DAPI, blue). Scale bar, 10 µm. **C**. Quantification of Keratin 8 phospho S34 fluorescence intensity at the cleavage furrow in control, 1 µM AZD1152 or 0.5 µM VX680-treated HeLa cells. Phospho S34 intensity at the cleavage furrow is divided by the cytoplasmic background. n=11 cells for control, n=9 cells for 1 µM AZD1152 treated and n=8 cells for 0.5 µM VX680 treated cells were quantified. One-way ANOVA with Dunnett’s post hoc test was performed. **** p<0.001. Data represent mean ± SEM (Standard Error Mean).

In addition to paraformaldehyde, we assessed Phospho S34 K8 localization after methanol fixation. These fixation methods differentially influence antibody epitope recognition: paraformaldehyde preserves both soluble and insoluble protein pools, whereas methanol fixation removes a substantial portion of soluble proteins, thereby enriching the detection of insoluble structures [32, 33]. As a result, phosphorylated K8 (pS34 K8) staining is less prominent in methanol-fixed cells compared to paraformaldehyde-fixed cells, consistent with its predominant association with the soluble fraction. Yet, during the early stages of furrow formation, phospho S34 K8 remains detectable at the cleavage furrow, suggesting that it is initially associated with the insoluble keratin pool, but gradually shifts toward the soluble fraction as the furrow progresses and regresses **(Supplementary Figure S5).**

### Keratin 8 and Aurora B kinase interact in a cell cycle-dependent manner

The findings above suggest temporal and spatial regulation of Keratin 8 phosphorylation by Aurora B kinase. To elaborate on the interaction between Keratin 8 and Aurora B, we tested whether Aurora B physically associates with Keratin 8 during cell division. For this, we employed Aurora B-GFP-expressing cells where GFP-tagged Aurora B is expressed under its own promoter using bacterial artificial chromosome (BAC) transgenomics in HeLa cells [34]. We pulled down Aurora B-GFP from the cell extracts arrested in interphase and mitosis. Aurora B interacted with Keratin 8 in mitosis but not in interphase **(Figure 5A).**

**Figure 5.**
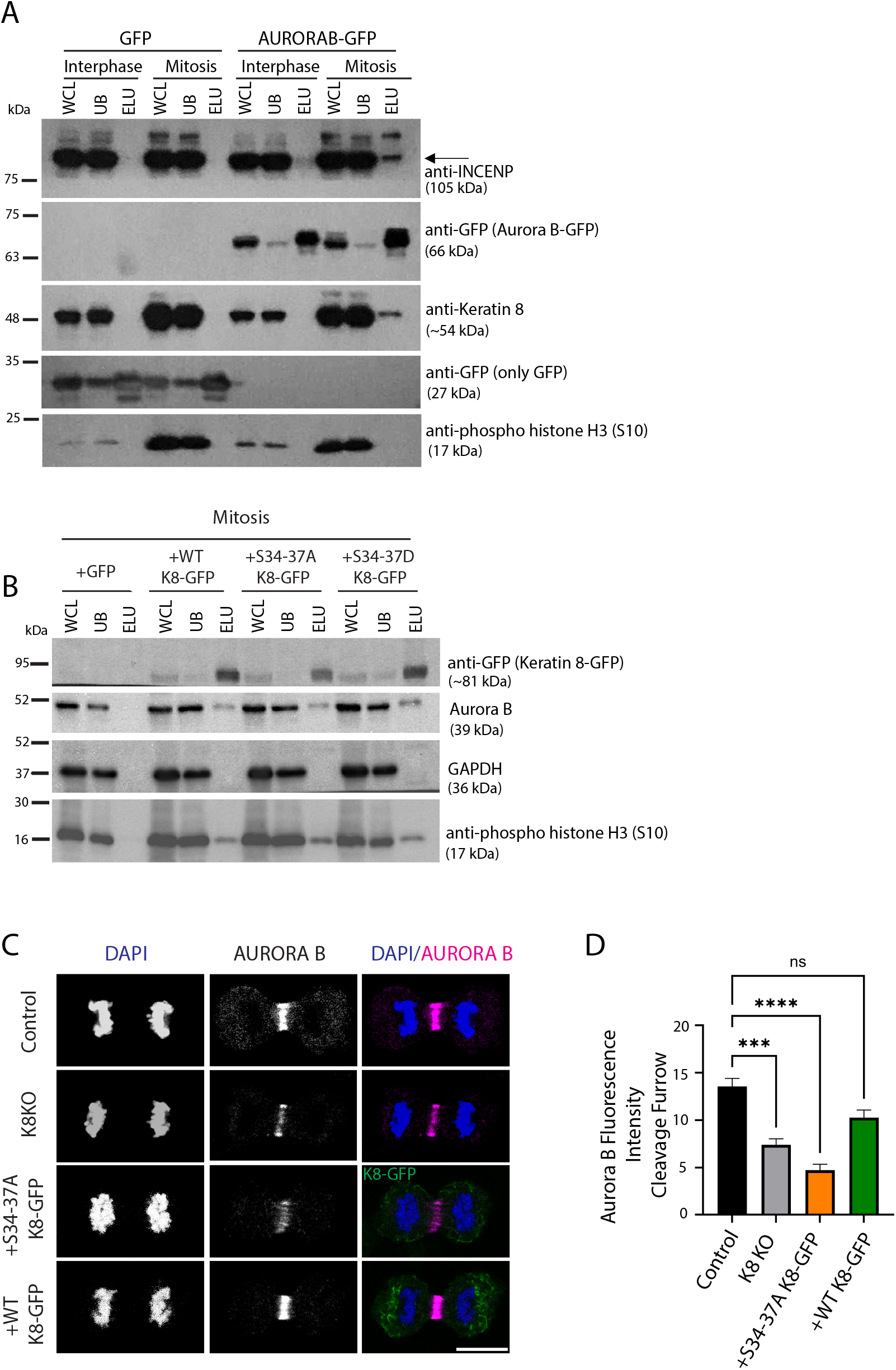
Keratin 8 interacts with Aurora B in a mitosis-dependent manner and facilitates its targeting to the midzone. **A**. GFP pulldown of HeLa cells stably expressing Aurora B-GFP in BAC (Bacterial Artificial Chromosome) (n=2). Immunoblotting of whole cell lysates (WCL), unbound (UB) and elutes (ELU) from control (Empty GFP) and Aurora B-GFP-expressing interphase and mitosis cells using anti-INCENP (expected band is indicated by an arrow), anti-Keratin 8 and anti-phospho histone H3 (S10). **B**. GFP pulldown of HeLa cells stably expressing Keratin 8-GFP (n=3). Western blotting analysis of whole cell lysates (WCL), unbound (UB) and elute (ELU) fractions obtained from K8 KO (Keratin 8 Knockout), HeLa cells expressing stable GFP (as control), WT K8-GFP, S34-37A K8-GFP and S34-37D K8-GFP. anti-GFP, anti-Aurora B, anti-phospho Histone H3(S10) and GAPDH antibodies were used. **C**. Representative images of Aurora B kinase localization in control (n=20), K8 KO (n=18), K8 KO HeLa cells expressing S34-37A K8-GFP(n=18) and WT K8-GFP (n=18) from three independent experiments. Anaphase cells are immunostained against Aurora B (magenta) and DNA (DAPI, blue). Scale bar, 10 µm. **D**. Quantification of the mean intensity of Aurora B localization at the midzone in control (n=20), K8 KO (Keratin 8 Knockout) (n=18), K8 KO HeLa cells expressing S34-37A K8-GFP (n=18), and WT K8-GFP (n=18) from three independent experiments. The Kruskal-Wallis test was performed. ****p<0.0001, *** p=0.0007, n.s., not significant. Data represent mean ± SEM.

In parallel, we performed GFP pulldown from WT K8-GFP and mutant (S34-37A and S34-37D) K8-GFP-expressing K8 KO cells. Empty GFP-expressing K8 KO cells were used as a control **(Figure 5B, Supplementary Figure S6-B)**. In agreement with the Aurora B GFP pulldown, the interaction between K8-GFP and Aurora B is pronounced during mitosis. Notably, all proteoforms, including both the phosphomutant and phosphomimetic forms of K8, significantly interacted with Aurora B during mitosis **(Figure 5B)**. This interaction was significantly reduced during interphase in the wild type and phosphomutant forms, but not in the phosphomimetic form **(Supplementary Figure S6A-B).**

To investigate the functional interaction between Keratin 8 and Aurora B kinase, we examined the subcellular localization of Aurora B kinase in the absence of K8 during cytokinesis using immunostaining in fixed cells. In control cells, Aurora B was localized at the spindle midzone; however, its localization was significantly reduced in K8 KO cells and cells expressing the S34-37A mutant. The expression of wild-type K8 was able to restore the spindle midzone localization of Aurora B in the K8 KO cells **(Figure 5C, D)**. Interchromatin distances of cells under different conditions were comparable, suggesting that the measurements were taken from equivalent stages of cell division **(Supplementary Figure S6C)**. Total Aurora B protein expression levels were unchanged in control, K8 KO and K8 KO + S34-37A K8-GFP-expressing cells **(Supplementary Figure S2C)**, indicating that the defect lies in Aurora B targeting rather than in its expression. Overall, these findings suggest a functional interaction between Keratin 8 and Aurora B kinase. Keratin 8 facilitates targeting of Aurora B kinase to the midzone spindle in keratin-expressing cells, which promotes localized keratin filament disassembly at the cleavage furrow.

## Discussion

Intermediate filaments are the least understood part of the cytoskeleton and have mostly been neglected by researchers studying cell division. Yet, it has long been recognized that intermediate filaments must be dissolved, either locally or globally, for cytokinesis to succeed. The regulation of intermediate filaments usually involves phosphorylation. Phosphorylation of nuclear lamins by CDK1 has long served as a paradigm for both intermediate filament and CDK1 biology [35, 36]. Aurora B-dependent phosphorylation of vimentin is also a classic example of the role of spatially regulated kinase activity in cell division [37]. However, very little is known about the cell cycle-dependent organization of Keratin filaments. Here, by combining mass spectrometry and microscopy, we showed that cytokinesis-specific and Aurora B-dependent phosphorylation of Keratin 8 decorates the cleavage furrow and midzone spindle, and it triggers the disintegration of Keratin bundles at the cleavage furrow.

Keratin filament reorganization is known to be tightly regulated by post-translational modifications. Phosphorylation, in particular, increases keratin solubility and promotes the disassembly of the keratin network [11, 38]. Previous studies have shown that site-specific phosphorylation solubilizes keratin filaments and disrupts network architecture [31, 39, 40]. By employing Aurora kinase inhibitors, Field et al. showed that the disassembly of keratin filaments between sister asters in *Xenopus egg* extracts requires Aurora B kinase activity [28]. However, that study did not identify any phosphosites or test the role of non-phosphorylatable mutant keratin. In this study, we mapped six Aurora B-dependent phosphorylation sites in Keratin 8, focusing on Serine 34, pS34 K8. Aurora B kinase phosphorylates K8 S34 at the onset of anaphase as a cleavage furrow begins to form. pS34 K8 decorates the cleavage furrow and spindle midzone and persists until furrow ingression is complete. The confined localization of pS34-K8 to the furrow and midzone suggests that keratin solubility is spatially regulated during cell division. We propose that Aurora B-mediated phosphorylation promotes local disassembly of keratin filaments specifically at the cleavage furrow. Similar to *Xenopus egg* extract [28], in human cells, we observed that Aurora B inhibition or expression of non-phosphorylatable K8 leads to persistent keratin bundles at the cleavage furrow and results in cytokinesis defects. Acute depletion of K8 or the introduction of additional mutations within other detected Aurora B phosphorylation sites on K8 will likely exacerbate the multinucleation phenotype observed in HeLa cells.

Previous work on the inhibition of cytokinesis by Aurora B inhibitors assumed that these drugs work by blocking signaling pathways leading to RhoA activation at the furrow cortex [18, 41]. Our results show that an additional factor is the blockade of furrow ingression by keratin filaments when they cannot be phosphorylated by Aurora B. Importantly, we observed that Aurora B kinase and Keratin 8 interact during mitosis and cytokinesis. The necessity of Aurora B-dependent Keratin phosphorylation for the ingression of the cleavage furrow and Keratin 8’s role in targeting Aurora B to the chromosomes and midzone suggest their functional interaction.

The phospho S34 Keratin antibody generated in this study provides a readout of the spatial extent of Aurora B kinase activity as well as a biomarker for cytokinesis. Spatial regulation of Aurora B kinase was previously demonstrated using a live cell Förster Resonance Energy Transfer (FRET) biosensor [42, 43], which has the advantage of temporal readout, but our antibody is technically simpler to implement and useful for fixed samples. Multiple anti-mitotic phospho-antibodies are used to detect mitotic cells including phospho Histone H3 antibodies [44], however, it is not feasible to distinguish between mitosis and cytokinesis cells with those [45]. Phospho S34 Keratin would serve as a great biochemical biomarker to examine cytokinesis cells in the population. Given the fact that Keratin 8 is a widely used tumor diagnostic biomarker in epithelial malignancies [3], anti-phospho S34 Keratin antibodies would have great value in assessing cell cycle progression in cancer cells and monitoring the response of anti-mitotic cancer chemotherapy.

The Keratin code refers to the cell-specific combination and modification of keratin proteins that shape the structure and function of the epithelial cytoskeleton, enabling diverse and adaptable epithelial behaviors [46]. Our findings extend this concept to cell cycle–specific keratin forms, demonstrating that keratin structure can be dynamically controlled through temporally and spatially restricted post-translational modifications. It will be important to determine the “readers and writers” [47] of different phospho-proteoforms of Keratins at different cell cycle stages to investigate their interactions and functional relevance.

Our findings extend the paradigm of cytoskeletal regulation during mitosis by demonstrating that epithelial keratins, long viewed mainly as structural scaffolds and diagnostic markers, are active participants in cytokinesis through Aurora B-mediated phosphorylation. While previous studies primarily focused on Aurora B’s role in regulating actomyosin contractility and RhoA signaling, our work uncovers a parallel pathway in which Aurora B locally disassembles keratin filaments to permit furrow ingression. This mechanistic insight is especially relevant in cancer biology, as keratin 8 is abundantly expressed in epithelial malignancies, where uncontrolled proliferation requires the completion of cell division. K8/K18 have been implicated in tumorigenesis and multiple oncogenic signaling pathways. K8 knockdown in A431 cells leads to reduced tumorigenic behavior both *in vitro* and in vivo [48]. This effect is likely due to the involvement of K8/K18 in the regulation of multiple oncogenic signaling pathways. K8/K18 expression stabilizes Akt activity to promote cell survival, and loss of keratin expression reduces Akt activity and induces apoptosis [49]. In hepatoma cells, K8/K18 influence adhesion and migration properties by modulating integrin/FAK/PKC-dependent signalings [50]. The interaction of K8/18 with 14-3-3 proteins regulates phosphorylation-dependent binding of 14-3-3 to diverse signaling molecules, including Raf-1 kinase, Bad, and cdc25 phosphatase [51]. Keratin-bound Raf is sequestered in an inactive state, but upon oxidative or toxin-induced stress, Raf hyperphosphorylation disrupts its association with K8/18, leading to its release and activation, in parallel with enhanced 14-3-3 binding to phosphorylated K18. Thus K8/18 act as dynamic interplayers to modulate Raf signaling during cell stress [52]. Moreover, keratins are direct substrates of the stress kinase p38 [53] and cAMP-dependent protein kinase, protein kinase C [54], further emphasizing their role as regulatory hubs in kinase networks. Our analysis contributes to the expanding understanding of the keratin-dependent regulatory network by revealing its connection to the master mitotic kinase, Aurora B.

Blocking mitosis is a longstanding strategy in anti-cancer drug development, but it has been difficult to selectively target cell division in cancer cells. For example, potent and specific inhibitors of Aurora kinase that were effective in mouse tumor models failed in clinical trials because they blocked cell division in white blood cell progenitors in the bone marrow [55, 56]. Unlike Aurora B kinase, Keratin 8 is only expressed in epithelial cells; therefore, their functional dependency should be cell type and differentiation-dependent. In principle, a drug that targets Keratin 8 disassembly by Aurora B kinase should have much higher selectivity for carcinoma treatment than a pan Aurora inhibitor. More work is required to decipher the molecular details of Keratin keratin-dependent function of Aurora B and downstream Aurora B-dependent keratin 8 disassembly, but in principle, these processes should be druggable. Furthermore, since K8/K18 integrate diverse oncogenic pathways to mediate survival and migration, their regulation by Aurora B may represent a critical point where keratin remodeling interfaces with cancer-driving kinase cascades in epithelial malignancies.

This study, for the first time, provided a molecular understanding of Aurora B kinase and Keratin 8 interaction and identified Keratin phosphorylation sites that are phosphorylated by Aurora B kinase in a short time window at a confined region, which is indispensable for cytokinesis. The major limitations of this study are the reliance on cancer-derived cell lines, which may not fully recapitulate the physiological regulation of tissue architecture. Future studies characterizing the anti–pS34-K8 antibody in tumor samples and assessing the functional relevance of the identified phosphorylation sites in animal models will be important to extend these findings. Notably, large-scale proteomic analyses of cancer tissues have reported deregulation of the phosphorylation sites characterized in this study [57, 58]. Phosphorylation of K8 at serine 37 was significantly increased in clear cell renal carcinoma tissues [58]. Future studies on Aurora B-Keratin interaction in epithelial malignancies will be important to determine its relevance for cancer progression and therapeutic targeting.

Overall, our study demonstrates a functional interaction between Aurora B and Keratin 8 for successful cytokinesis in Keratin-expressing cells, which provides insight into the mechanism of cell-type/cancer specific pathways during cytokinesis.

## Materials and Methods

### Antibodies

The following antibodies were used; mouse monoclonal α-tubulin DM1A (Cell Signaling 3873S, Western blotting 1:1000, immunofluorescence microscopy 1:1000), mouse monoclonal Keratin 8 (Santa Cruz, sc-8020, Western blotting 1:1000, immunofluorescence microscopy 1:1000), rabbit monoclonal Aurora B (Abcam, ab2254, Western blotting 1:1500) mouse monoclonal GFP (Roche, 11814460001, Western blotting 1: 1000), rabbit polyclonal INCENP (Bethyl Laboratories, IHC-00060, Immunofluorescence microscopy 1:250), mouse monoclonal GAPDH (Cell Signaling, 97166S, Western blotting 1:4000), rabbit anti-Phospho Histone H3 (Santa Cruz, sc-8656-R, Western blotting 1:200). For generating antibodies against phospho S34 Keratin 8 (produced by Davids Biotechnology), rabbits were immunized with the Keyhole Limpet Hemocyanin conjugated phospho-peptide five times. After antibody production, the serum is affinity-purified with the phospho-peptides. To get rid of antibodies recognizing the non-phosphopeptides, the affinity-purified serum is depleted with a non-phosphopeptide depletion matrix.

### Plasmids and Cloning

GFP-tagged WT Keratin 8 in pEGFP-N3 was kindly provided by Dr. Milind Vaidya (ACTREC). S34-35A, S35-36A, S34-37A, S34-35D and S34-37D constructs were cloned into the pEGFP-N3 vector by site-directed mutagenesis using WT Keratin 8-GFP. WT Keratin 8 was cloned into pGEX-6p-1 (WT-K8-GST) for in vitro assays.

### Cell Culture and Synchronization

HeLa S3(RRID: CVCL_0058) and MCF7 (RRID: CVCL_0031) cells were obtained from the American Type Culture Collection and routinely cultured in Dulbecco’s modified Eagle’s medium (DMEM) (Gibco, 41966029) containing 10 % Fetal Bovine Serum, 100 units/ml Penicillin and 100 μg/ml Streptomycin (Thermo,15140122) at 37°C and 5% CO_2_. To synchronize the cells in interphase, cells were incubated with DMEM including 2 mM Thymidine (Santa Cruz, 296542A and Calbiochem 6060-5GM) for 20 hours, then released into fresh DMEM for 8 hours. Cells were incubated with 2 mM Thymidine for another 17 hours. To arrest cells at prometaphase, cells were incubated in DMEM including 30 nM Nocodazole (Calbiochem, 487928) for 5 hours. Cells were harvested or fixed at the end of one hour after release from nocodazole to collect cytokinesis cells. To induce Aurora B inhibition at cytokinesis, cells were treated with 1 μM AZD1152 or 0.5 μM VX680 Aurora B inhibitors for 30 minutes. All experiments were performed using mycoplasma-free cells, which were routinely monitored by microscopic examination and PCR-based detection.

### CRISPR/Cas9 mediated Keratin 8 Knockout

Targeting guide with sequence 5’ CGAGGAGCTGATGCGGGAA 3’ and non-targeting guide with sequence 5’-ACGGAGGCTAAGCGTCGCAA-3’ were cloned into pLenti CRISPRv2 using BsmBI restriction digestion of the backbone using the protocol from Zhang lab. HeLa S3 cells were transiently transfected with the plentiCRISPRv2 plasmid containing the Keratin 8 guide and a non-targeting control by using Lipofectamine. Transfected cells were selected with puromycin. Serial dilution was used to prepare monoclonal cells. Loss of Keratin 8 expression in HeLa cells was detected by using immunoblotting.

### Pulldown of Aurora B-GFP

Aurora B-GFP-expressing cells were grown in 40% confluency and synchronized in interphase by double Thymidine Block and in mitosis by using S-trityl-L-cysteine (Sigma-Aldrich) [59]. Cell pellets were lysed in ice-cold lysis buffer (20mM Tris-CL pH7.4, 150mM NaCl, 1mM MgCl2, 10%Glycerol, 0.5mM EDTA, 10mM NaF, 0.5%NP-40, 1mM beta-glycerolphosphate, 1mM sodium pyrophosphate, 1mM sodium orthovanadate, 1mM DTT, EDTA-free protease inhibitor (Pierce, 88266) by pipetting and passing through a 25-gauge needle. Cell lysates were centrifuged at 14000 rpm for 15 min at +4°C. Dilution buffer (20mM Tris-CL pH7.4, 150mM NaCl, 1mM MgCl2, 10%Glycerol, 0.5mM EDTA, 10mM NaF) was added to the supernatant in 2:3 proportions. GFP-Trap®_A (Chromotek, gta-20) was used for pulling down Aurora B-GFP. The diluted lysate was mixed with the beads and rotated for 3 hours at 4°C. After washing the beads with dilution buffer, for elution, beads were re-suspended in 2X Laemmli Sample Buffer (4% (w/v) SDS, 20% Glycerol, 120mM Tris-Cl (pH 6.8), 0.02% (w/v) bromophenol blue, 100mM DTT) and boiled for 10 min at 95°C and SDS-PAGE and Western Blotting were performed.

### Pulldown of K8-GFP

K8-GFP-expressing cells were grown in 35-40 % confluency and synchronized in interphase by double thymidine block and in mitosis by using Nocodazole (Calbiochem,487928)) as mentioned above. Cell pellets were lysed in ice cold lysis buffer (20 mM Tris/cl pH7.4, 150 mM NaCl, 10 Mm NaF, 0.5%NP-40, 1 mM MgCl2, 0.5 Mm EDTA, 10% glycerol, 1mM DTT, 200 Mm PMSF (MP, 195381), ¼ protease inhibitor mini tablets (EDTA free, Thermo, 88666), PhosSTOP (Roche)) by pipetting and passing through 25-gauge needle. Cell lysates were centrifuged at 14000 g for 10 min at 4°C. Dilution buffer (20 mM Tris/cl pH7.4, 150 mM NaCl, 10 Mm NaF, 1 mM MgCl2, 0.5 Mm EDTA, 10% glycerol, 1mM DTT, 200 Mm PMSF (MP, 195381), ¼ protease inhibitor mini tablets (EDTA free, Thermo, 88666), ¼ PhosSTOP (Roche)) was added to the supernatant in 2:3 proportions. GFP-Trap®_Agarose (Chromotek, gta-100) was used for pulling down K8-GFP. GFP-Trap®_Agarose (Chromotek, gta-100) beads were pre-blocked with 2% BSA in lysis buffer at +4°C for 1-2 h or overnight. GFP beads were washed with ice-cold dilution before use. The diluted lysate was mixed with the beads and rotated for one hour at 4°C. After washing the beads with dilution buffer, for elution, beads were re-suspended in 2X Laemmli Sample Buffer (4% (w/v) SDS, 20% Glycerol, 120mM Tris-Cl (pH 6.8), 0.02% (w/v) bromophenol blue, 100mM DTT) and boiled for 5 min at 95°C and SDS-PAGE and Western Blotting were performed.

### Transfection

WT and mutant K8-GFP constructs were transfected into HeLa S3 cells with Lipofectamine 2000 by following the manufacturer’s instructions for immunostaining and live-cell imaging experiments. HeLa S3 cells were transfected with vectors 8-24 hours before the thymidine block. 300-1500 ng DNA was incubated with 1-5 μl Lipofectamine 2000 in 150 μl Optimem (Thermo Fisher, 31985047) for 10 minutes at room temperature and then given to the cells. The media was changed 8-10 hours after transfection.

### Immunofluorescence

Cells were grown on coverslips and fixed with 3% paraformaldehyde for 15 min at 37°C or with methanol at −20°C for 10 min. Fixed coverslips were washed with PBS-0.1% Triton-X (PBS-Tx) and incubated with blocking solution (2% BSA in PBS-Tx) overnight at 4°C or for 30 minutes at room temperature. Incubation of the primary antibody in 2% BSA in PBS-Tx was done at 4°C overnight or at room temperature for 2 hours. Secondary antibody in 2% BSA in PBS-Tx was incubated at room temperature for 1 hour. Cell nuclei staining was done with 1 μg/ml DAPI in 2% BSA, PBS-Tx for 10 minutes at room temperature.

### Microscopy, Image Analysis and Statistical Analysis

Immunofluorescence intensities were quantified using ImageJ and FiJi software (Wayne Rasband, NIH). Briefly, RGB values of different spots at the midzone spindle or on the cleavage furrow were measured by keeping the area constant, and an average of these values was taken. The same process was applied to random spots on the cytoplasm of the cell, and the fluorescence intensities were normalized by dividing or subtracting the fluorescence intensity of the cytoplasm. For microscopy imaging, Leica DMi8/SP8 TCS-DLS, Leica DMi8 wide-field microscope using LAS X Software and NIS-Element Imaging software and Nikon Eclipse APO λ 100X/1.40 Oil objective lens were used. Graphs were plotted by GraphPad Prism 8.

### Recombinant protein expression and purification

The K8-GST vector was transformed into *Escherichia coli* BL21 competent cells by following the standard transformation protocol. A single colony was chosen and grown in 100 μg/ml Ampicillin LB at 37 °C until its OD600 reached 0.5. Then, bacteria were induced with 1 mM Isopropyl β-D-1-thiogalactopyranoside (IPTG). The recombinant protein was isolated from inclusion bodies. Briefly, bacteria were pelleted and lysed in a buffer containing 100 mM TrisCl pH 8, 5mM EDTA, 5mM DTT (Dithiothreitol) and 1X Protease Inhibitor (Roche, 11836170001) using a sonicator. The suspension was centrifuged at 13,500 rpm at 4°C for 1 hour. The pellet was re-suspended in a wash buffer (50 mM TrisCl pH 8, 10 mM EDTA, 5 mM DTT, 1 M NaCl, 1% NP40). After homogenization of the pellet, it was sonicated; this step was repeated one more time. After the washing steps, the pellet was extracted with extraction buffer (8 M Urea, 5 mM DTT, 2 mM EDTA, 10 mM TrisCl pH 8) and homogenized with a tissue grinder homogenizer. The suspension was pelleted again, and the supernatant was used for protein purification. To refold Keratin 8-GST, the supernatant was dialyzed with dialysis buffer (25 mM Hepes pH 7.4, 100 mM KCl, 5mM MgCl2, 0.5 mM EGTA). The Glutathione beads were incubated with the supernatant overnight. The beads were washed with a wash buffer (25 mM Hepes, 150 mM KCl, 5mM MgCl2, 0.5 mM EGTA, 1 mM DTT, 0.01% NP40) at 4°C twice. Harsh wash buffer (25 mM Hepes, 600 mM KCl, 5mM MgCl2, 0.5 mM EGTA, 1 mM DTT, 0.01% NP40) was used for the third wash. WT-GST-K8 was eluted in a buffer containing 100 mM TrisCl pH 8.0, fresh 10 mM Reduced Glutathione and 1mM DTT.

### *In vitro* kinase assay and Mass Spectrometry

Isolated WT-K8-GST was incubated with purified Aurora B complex, Aurora B and INCENP fragments were co-expressed from the bicistronic vector pGEX-2rbs with GST-tag at the N-terminus of Aurora B. The Aurora B complex was purified with GST sepharose and eluted from the beads by cleaving the GST-tag by PreScission protease in a buffer including 50 mM Tris pH 7.5, 150 mM NaCl, 1 mM EDTA, 1 mM DTT, 10 µg/ml leupeptin and 10 µg/ml pepstatin. The purified Aurora B complex was a gift from Dr. Masanori Mishima (University of Warwick). For the *in vitro* kinase assay, 25 µg WT-K8-GST and 0.75 µg Aurora B complex were incubated in 100 µl kinase reaction buffer including 20 mM PIPES pH 7, 2 mM MgCl_2_, 2 mM EGTA, 100 mM NaCl and 0.2 mM ATP at 30°C for 40 minutes. The kinase reaction was dissolved in 50 µl 3X SDS Blue Loading Buffer supplemented with 100 mM DTT by boiling it at 85°C for 10 minutes. After alkylation with 100 mM IA (I6125, Sigma Aldrich), the samples were loaded into 12% Tris-Glycine Precast Gels (Pierce, 25247) and stained with Page Blue Protein Staining (Thermo Fisher Scientific, 24620). The band corresponding to Keratin 8-GST (66-81 kDa) was cut and after the washing steps, the gel plugs were digested by using 1:50 (Trypsin: Protein amount ratio) Sequencing Grade Modified Trypsin (Promega) at 37 °C overnight. Digests were desalted by Stage Tipping using Empore C18 47mm disks and resuspended in 5% Formic Acid and 5% Acetonitrile for LC-MS/MS analysis.

### Mass Spectrometry Data Acquisitions and Processing

The peptides were subjected to a reversed-phase Nano LC-MS/MS (EASY-nLC, Thermo) connected to a Q Exactive quadrupole Orbitrap mass spectrometer (Thermo Fisher Scientific, Bremen). The peptides in the fractions were directly loaded onto an in-house packed 100 μm i.d. × 17 cm C18 column (Reprosil-Gold C18, 5 μm, 200Å, Dr. Maisch). Survey spectra were acquired on the Orbitrap with a resolution of 70,000 and for MS2 resolution 17,500. Raw data files were processed with Protein Discoverer (version 1.4 Thermo Scientific) and MaxQuant (version 1.5.2.8) for protein identification. The raw data were searched against a database including the Keratin 8-GST sequence. For the PRM analysis, an inclusion lists were prepared and the samples were analysed with a 60 min linear gradient, the following MS parameters: Dynamic exclusion 10 ppm, MS2 resolution 35.000, AGC target 2e5, isolation window 1.2 m/z, nce 28. The transition lists were created in Skyline v3.1 software (MacCoss Lab) with the following transition settings: precursor charges 2,3,4,5; ion charges 1,2 and eight product ions. MS and MS/MS mass accuracy were set to 10 ppm.

## Abbreviations

BAC: Bacterial Artificial Chromosome
BSA: Bovine Serum Albumin
CDK1: Cyclin-Dependent Kinase 1
CPC: Chromosome Passenger Complex
DMEM: Dulbecco’s Modified Eagle’s Medium
DTT: Dithiothreitol
EDTA: Ethylenediaminetetraacetic Acid
EGTA: Ethylene glycol-bis (β-aminoethyl ether)-N,N,N’,N’-tetraacetic Acid
GFP: Green Fluorescent Protein
K8: Keratin 8
K8 KO: Keratin 8 Knockout
K18: Keratin 18
NP40: Nonidet P-40 (Nonyl Phenoxypolyethoxylethanol)
PBS-Tx: Phosphate-Buffered Saline + Triton X-100
PMSF: Phenylmethylsulfonyl Fluoride
PRM: Parallel Reaction Monitoring
SDS-PAGE: Sodium Dodecyl Sulfate–Polyacrylamide Gel Electrophoresis

## Author Contributions

NO and TJM conceived the project; NO designed/supervised experiments. BH, MHQ and HA prepared biological samples; BH and HA performed cell biology, molecular cloning, biochemistry and imaging experiments; analyzed the data and prepared the figures. HB, HA, XW analyzed microscopy images and performed statistical analysis. VB, AIN and NO performed Mass Spectrometry and PRM experiments and analyzed data. BH, NO, TJM wrote the original draft; NO reviewed/edited the manuscript with input from all authors.

## Acknowledgement

We thank Dr. Milind Vaidya from ACTRECT for providing WT K8-GFP vector. We thank Dr. Masanori Mishima from Warwick University for supplying purified active Aurora B; Dattatreya Mellacheruvu from the Department of Pathology, Michigan University, for the help with the PRM experiment; Gürkan Mollaoğlu, M. Göksu Özlü, Öykü Kaya, Merve Yiğin and Ceren Evcil for their help in the experiments. Artür Manukyan from Max Delbrück Center for his help with statistical analysis. BH was supported by a scholarship from the TUBITAK-2211E program. This work is also supported by TUBITAK (118Z832) and International Center for Genetic Engineering and Biotechnology (ICGEB) (CRP/23/011) awarded to N.O. The graphical abstract was created in BioRender. Ozlu, N. (2025) https://BioRender.com/wcnpygh

## Data Availability

The main body of data, including analyses and images, is available in the article or its supplementary information. Source of data are available from the corresponding author upon reasonable request.

## Supplementary Information

Figure S1. Mapping of Aurora B-dependent phosphorylation of Keratin 8 in vitro.

Figure S2. Non-phosphorylatable Keratin 8 mutation causes multinucleation.

Figure S3. Phospho S34 Keratin 8 antibody is not detected in Keratin 8 knockout HeLa cells.

Figure S4. Keratin 8 S34 phosphorylation localizes specifically to the cleavage furrow in MCF7 cells in an Aurora B-dependent manner.

Figure S5. Keratin 8 S34 phosphorylation at the cleavage furrow is more pronounced in paraformaldehyde-fixed cells than in methanol-fixed cells.

Figure S6. Keratin 8 is associated with Aurora B in mitosis-dependent manner.

Video S1. Live imaging video of control (DMSO-treated) Keratin 8 knockout (K8 KO) HeLa cells expressing K8-GFP during cytokinesis.

Video S2. Live imaging video of Aurora inhibitor (AZD1152) treated K8 KO HeLa cells expressing K8-GFP during cytokinesis.

Video S3. Live imaging video of Aurora inhibitor (VX680) treated K8 KO HeLa cells expressing K8-GFP during cytokinesis.

Video S4. Live imaging video of K8 KO HeLa cells expressing WT K8-GFP and H2B-mCherry during cell division. K8-GFP and H2B-mCherry are merged.

Video S5. Live imaging video of K8 KO HeLa cells expressing S34-37A K8-GFP and H2B-mCherry during cell division. K8-GFP and H2B-mCherry are merged.

Video S6. Live imaging video of K8 KO HeLa cells expressing S34-35D K8-GFP and H2B-mCherry during cell division. K8-GFP and H2B-mCherry are merged.

