## Supplementary Figures S1_S6 for "Spatial control of Keratin 8 phosphorylation by Aurora B facilitates cytokinesis in cancer cells of epithelial origin"

**SUPPORTING INFORMATION**

**Description of Additional Files:**

**Supplementary Figures (this document): 6**
**Supplementary Videos: 6**

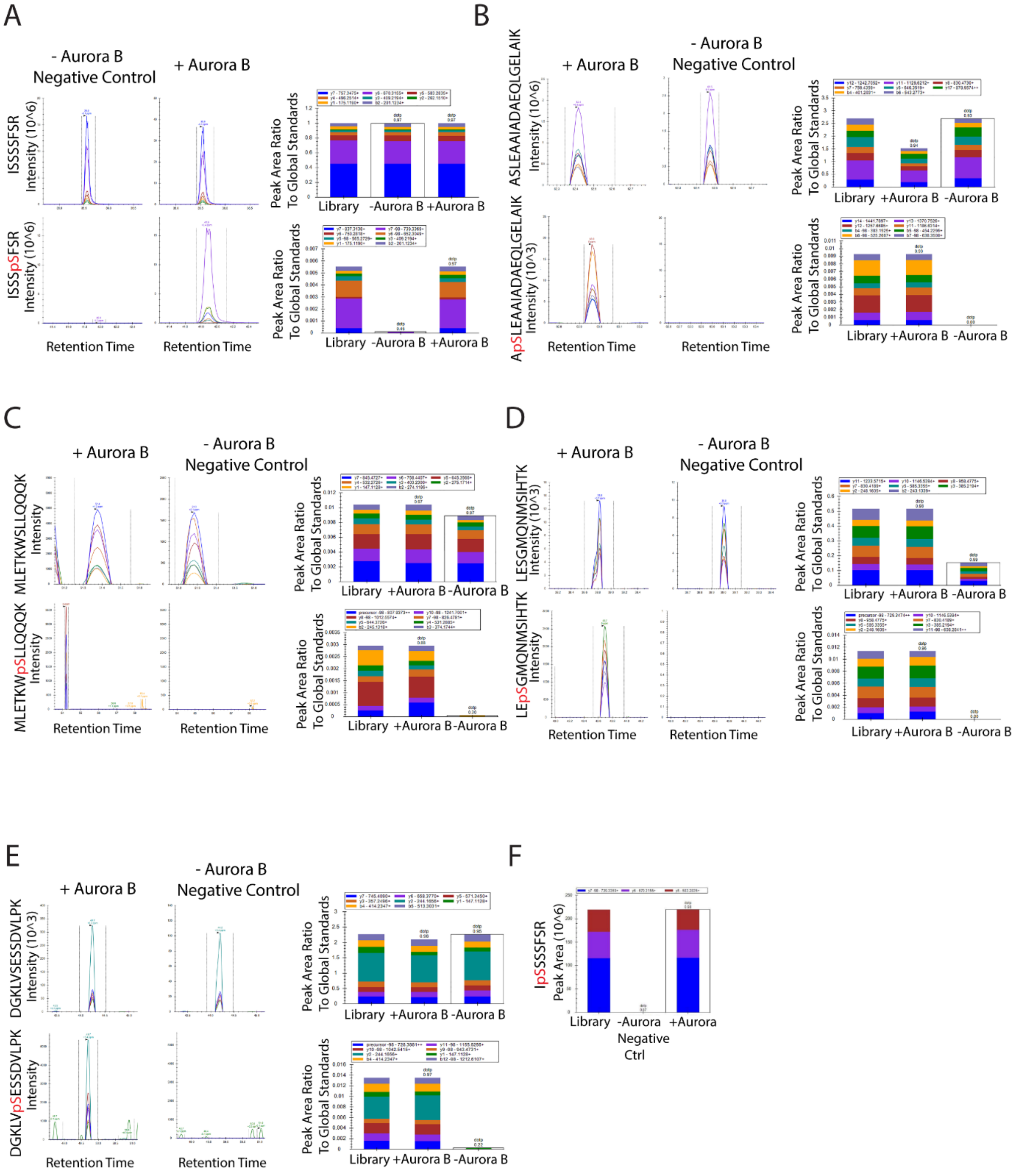


**Figure S1. Mapping of Aurora B-dependent phosphorylation of Keratin 8 *in vitro.***

Parallel Reaction Monitoring-Mass Spectrometry (PRM-MS) analysis of *in vitro* phosphorylation peptides of Keratin 8 by Aurora B (n=1). The peak area quantification of the non phospho and phospho peptides of Keratin 8 with Aurora B (+Aurora B) and without Aurora B (-Aurora B) conditions. The transitions of the non-phospho or phospho peptide of K8 (without Aurora B (-Aurora B) and with Aurora B (+ Aurora B) (left) and their quantification (right) are shown for each peptide using Skyline. Each graph shows one peptide and the colored peaks show the transitions of the non -phosphopeptide (top) and phosphopeptide (bottom). The histogram on the right shows the total intensity of each transition (in different colors), for each transition list including m/z values.

**A.** non-phospho ISSSSFSR and phospho ISSSpSFSR (pS37) Keratin 8

**B.** non-phospho ASLEAAIADAEQLGELAIK and phospho ApSLEAAIADAEQLGELAIK (pS330) Keratin 8

**C.** non-phospho MLETKWSLLQQQK and phospho MLETKWpSLLQQQK (pS124) Keratin 8

**D.** non-phospho LESGMQNMSIHTK and phospho LEpSGMQNMSIHTK (pS404) Keratin 8

**E.** non-phospho DGKLVSESSDVLPK and DGKLVpSESSDVLPK (pS475) Keratin 8

**F.** Skyline analysis compares transitions of non-phospho ISSSSFSR and phospho IpSSSSFSR (pS34) peptide of K8 with the known library from in vitro assay (without Aurora B (-Aurora B) and with Aurora B (+ Aurora B)).

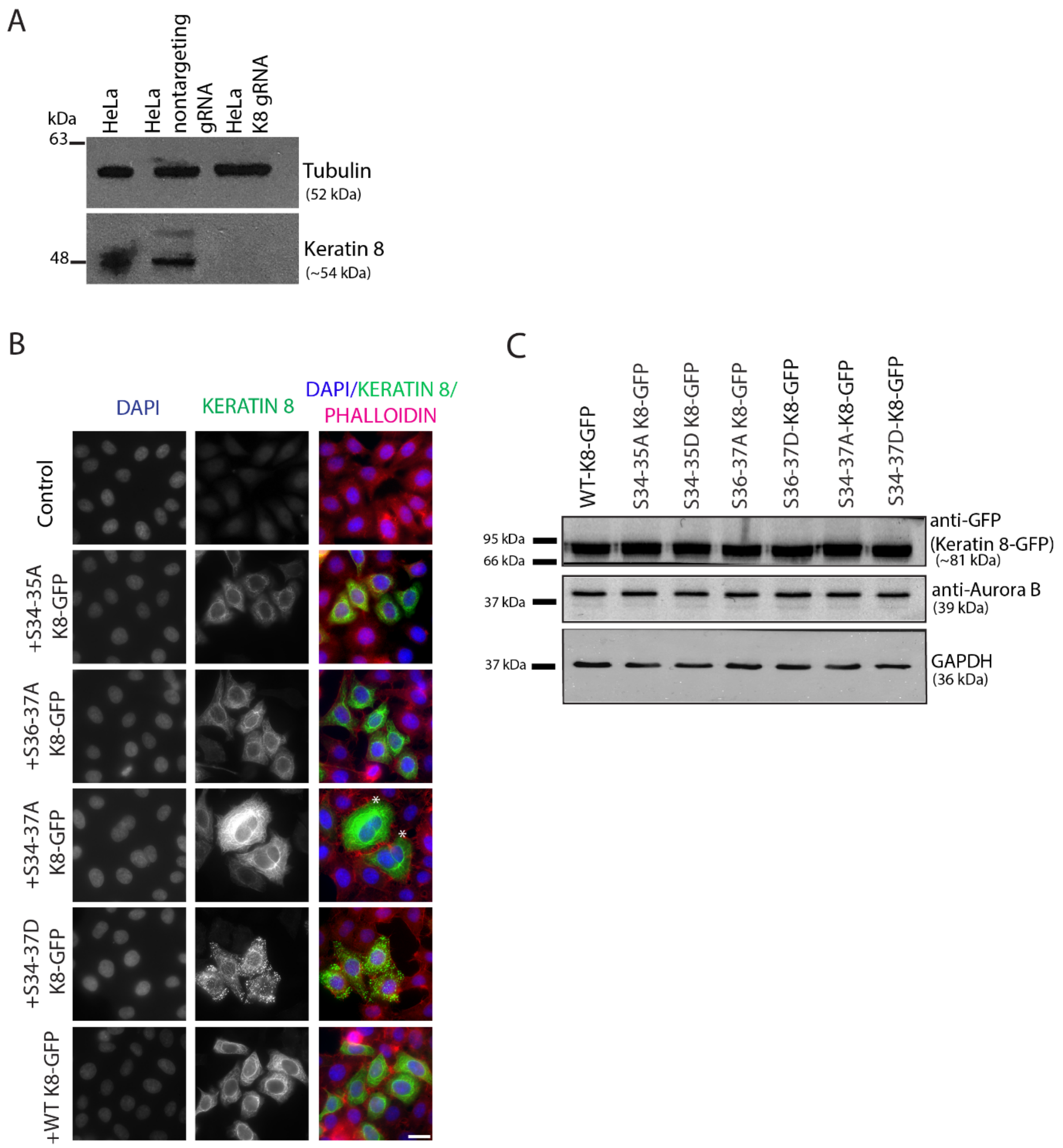


**Figure S2. Non-phosphorylatable Keratin 8 mutation causes multinucleation.**

**A.** Western Blotting analysis of Keratin 8 Knockout by detecting Keratin 8 and Tubulin protein levels in HeLa cells treated with nontargeting gRNA or K8 gRNA (n=3). Whole cell lysates were immunoblotted with anti-Keratin 8 and anti-α-Tubulin antibodies.

**B.** Imaging of fixed control (nontargeting gRNA) (n=3163) and Keratin 8 Knockout (K8 KO) HeLa cells expressing S34-35A K8-GFP (n=1568), S36-37A K8-GFP (n=796), S34-37A K8-GFP (n=1450), S34-37D K8-GFP(n=1371), and WT K8-GFP (n=1477). Cells were stained using phalloidin (magenta), K8-GFP (green), and DAPI (blue). Asterisks highlight multinucleated cells. Scale bar, 20 µm.

**C.** Western Blotting analysis in K8 KO HeLa cells expressing WT K8-GFP, S34-35A K8-GFP, S34-35D K8-GFP, S36-37A K8-GFP, S36-37D K8-GFP, S34-37A K8-GFP, and S34-37D K8-GFP (n=3). Whole cell lysates were immunoblotted with anti-GFP, anti-Aurora B, and GAPDH antibodies.

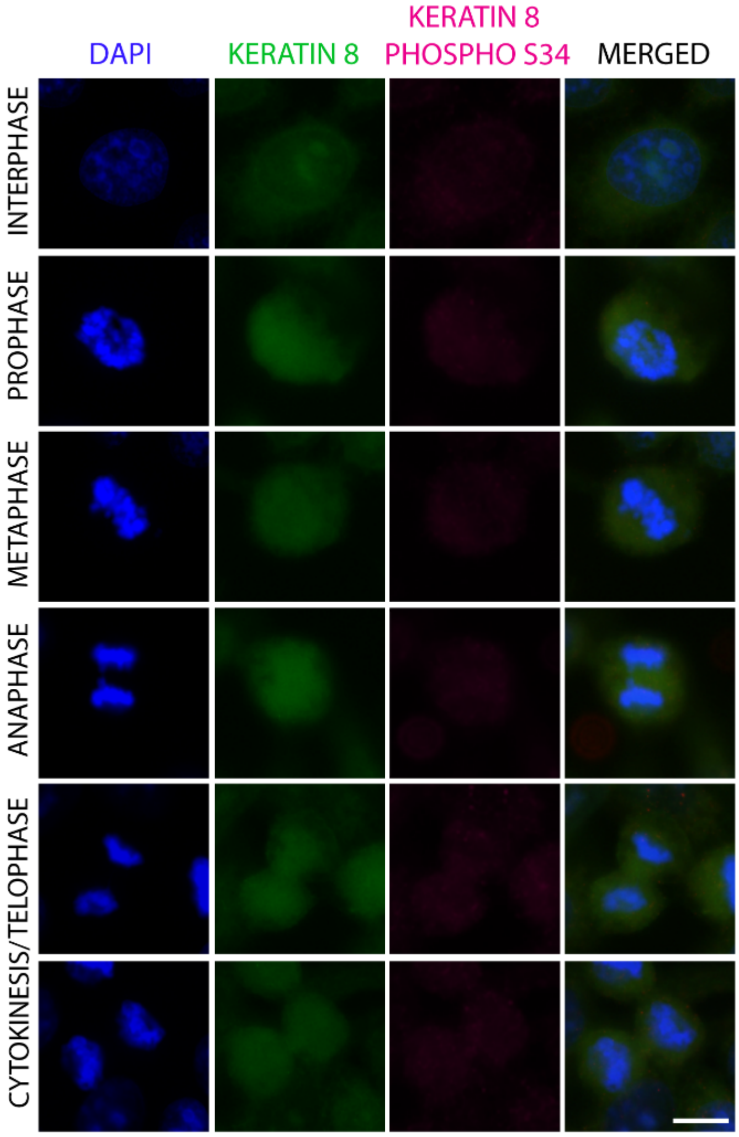


**Figure S3. Phospho S34 Keratin 8 antibody is not detected in Keratin 8 knockout HeLa cells**

Immunofluorescence staining of Keratin 8 Knockout HeLa cells at different cell cycle stages (interphase-cytokinesis) using Phospho S34 Keratin 8 (magenta) and Keratin 8 (green) antibodies and DAPI (blue) (n=2). Scale bar, 10 µm.

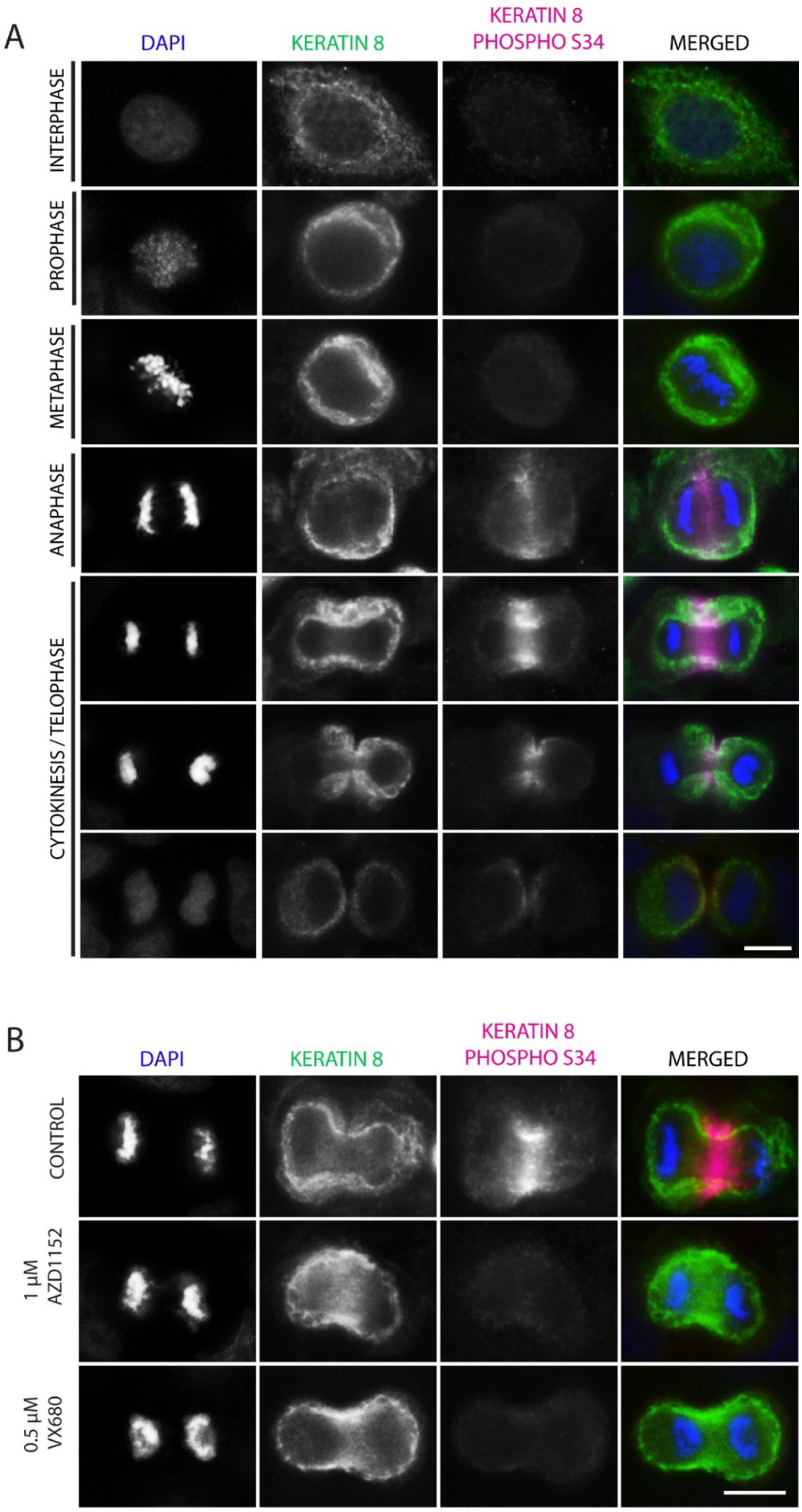


**Figure S4. Keratin 8 S34 phosphorylation localizes specifically to the cleavage furrow in MCF7 cells in an Aurora B-dependent manner.**

**A.** Immunofluorescence staining of MCF7 cells at different cell cycle stages (interphase-cytokinesis) using Keratin 8 (green), Keratin 8 phospho S34 (magenta) antibodies, and DAPI (blue) (n=3).

**B.** Immunofluorescence staining of control (DMSO) and Aurora B kinase inhibitor (1 µM AZD1152 or 0.5 µM VX680) treated MCF7 cells using Keratin 8 (green), Keratin 8 phospho S34 (magenta) antibodies, and DAPI (blue) (n=3). Scale bars, 10 µm.

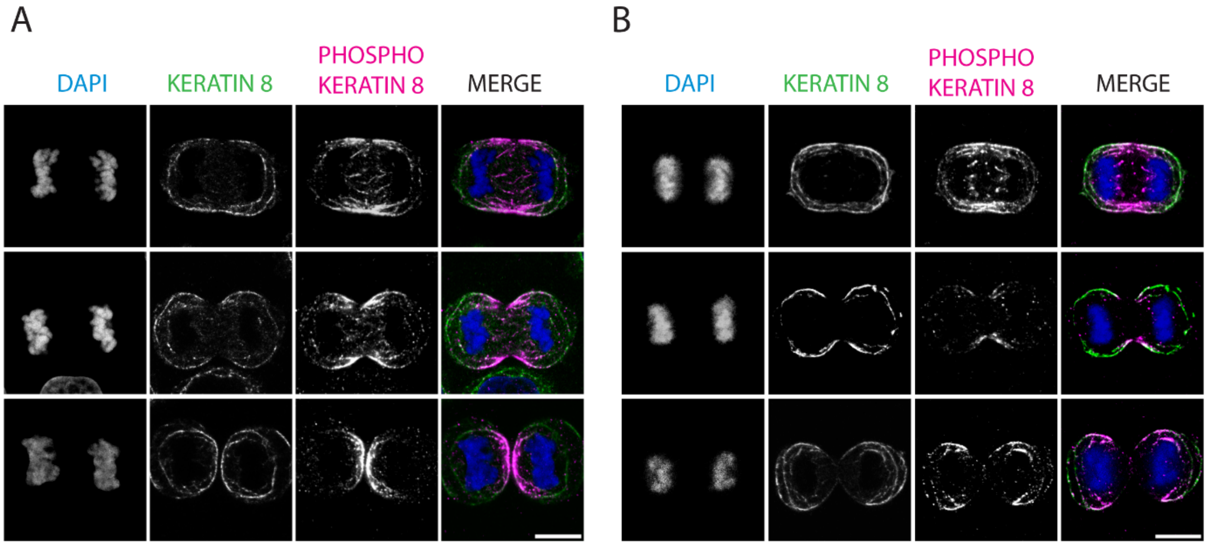


**Figure S5. Keratin 8 S34 phosphorylation at the cleavage furrow is more pronounced in paraformaldehyde-fixed cells than in methanol-fixed cells.**

Immunofluorescence staining of HeLa cells undergoing cytokinesis, labeled with K8-GFP (green), anti-phospho-Keratin 8 (Ser34) antibody (magenta), and DAPI (blue) (n=3). **A.** Cells were fixed with paraformaldehyde. **B.** Cells were fixed with methanol. Scale bars, 10 µm.

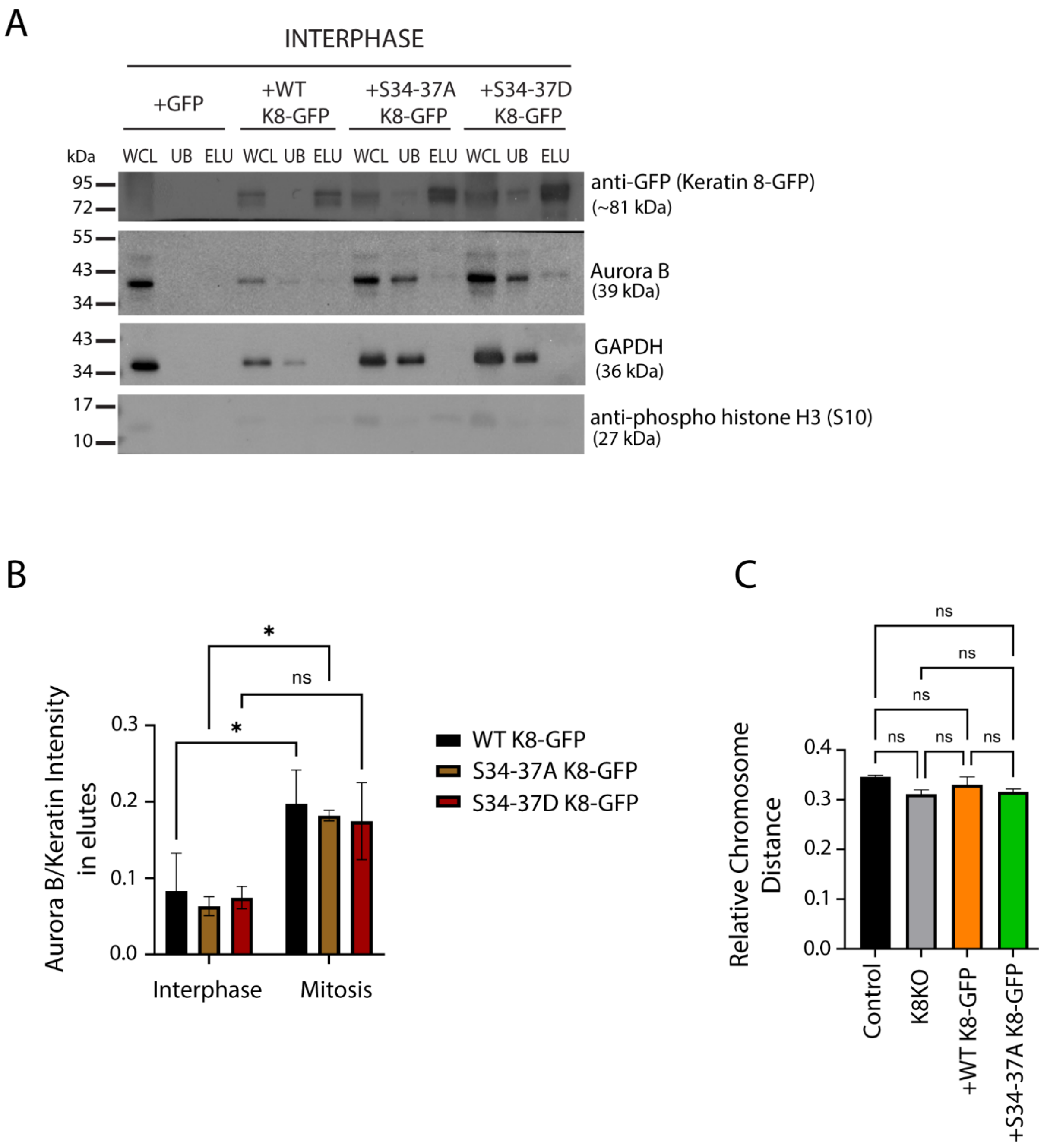


**Figure S6. Keratin 8 is associated with Aurora B in a mitosis-dependent manner.**

**A.** GFP pulldown of HeLa cells stably expressing K8-GFP synchronized in interphase (n=3). Western blotting analysis of whole cell lysates (WCL), unbound (UB), and elute (ELU) fractions obtained from Keratin 8 Knockout (KO) HeLa cells expressing GFP alone (as control), WT K8-GFP, S34-37A K8-GFP, or S34-37D K8-GFP. Blots were probed with anti-GFP, anti-Aurora B, anti-phospho Histone H3 (S10), and GAPDH antibodies.

**B.** Quantification of Aurora B co-elution in K8-GFP pulldown assays performed in interphase and mitosis using K8 Knockout HeLa cells expressing WT K8-GFP, S34-37A K8-GFP, and S34-37D K8-GFP (n=3). Aurora B levels were normalized to K8-GFP. Statistical analysis was performed using two-way ANOVA. *p = 0.0405; n.s., not significant. Data are presented as mean ± SEM.

**C.** Quantification of relative chromosome distance in Keratin 8 knockout HeLa cells expressing WT K8-GFP and S34-37A K8-GFP. Chromosome distances were normalized by whole cell length. 18 cells were analyzed in each of three independent experiments. Statistical analysis was performed using one-way ANOVA with Tukey's multiple-comparison test. n.s., not significant. Data are presented as mean ± SEM.

**Videos:**

**Video S1.** Live imaging video of control (DMSO-treated) Keratin 8 Knockout (K8 KO) HeLa cells expressing K8-GFP during cytokinesis.

**Video S2.** Live imaging video of Aurora inhibitor (AZD1152) treated K8 KO HeLa cells expressing K8-GFP during cytokinesis.

**Video S3.** Live imaging video of Aurora inhibitor (VX680) treated K8 KO HeLa cells expressing K8-GFP during cytokinesis.

**Video S4.** Live imaging video of K8 KO HeLa cells expressing WT K8-GFP and H2B-mCherry during cell division. K8-GFP and H2B-mCherry are merged.

**Video S5.** Live imaging video of K8 KO HeLa cells expressing S34-37A K8-GFP and H2B-mCherry during cell division. K8-GFP and H2B-mCherry are merged.

**Video S6.** Live imaging video of K8 KO HeLa cells expressing S34-35D K8-GFP and H2B-mCherry during cell division. K8-GFP and H2B-mCherry are merged.
